# The Fallacy and Bias of Averages on Vegetation Indices based Plant Phenotyping

**DOI:** 10.1101/2025.10.14.682309

**Authors:** Chun-Han Lee, Chih-Chen Tseng, Li-yu Daisy Liu, Tsu-Wei Chen

## Abstract

**Background:** Vegetation indices (VIs) from remote sensing are widely used for non-destructive plant phenotyping, often averaged across plots or image regions to represent each plot. However, according to Jensen’s inequality, which is known as the “fallacy of the average”, it can bias estimates when nonlinear relationships exist between VIs and target traits. To examine this issue, we systematically assessed the severity of this bias and tested a correction method. VI values were simulated using six beta distributions with varying shapes and skewness, and with normalized difference vegetation index (NDVI) images from a paddy rice experiment to evaluate bias under real conditions. Nonlinear link functions (concave, convex, logistic) with different noise levels were applied to model VI–trait relationships.

**Result:** The results showed that averaging under nonlinear relationships reduced predictive performance, lowering the coefficient of determination (R^2^) between true and predicted traits by up to 82%. In the rice NDVI simulation, R^2^ was reduced by up to 58% around the tillering stage. Our correction method, which predicts traits from VI before averaging, substantially mitigated bias, improving R^2^ by up to 0.68 depending on noise level, VI distribution, and link function. To facilitate application, we established an interactive R Shiny website enabling users to quantify potential biases and the efficacy of corrections within this workflow based on their own research conditions

**Conclusion:** In summary, averaging VIs without accounting for nonlinear relationships can introduce substantial bias and degrade phenotyping accuracy. This bias should be explicitly considered in phenotyping analyses, and correction methods applied when appropriate to improve reliability.

## Background

With the advances in various remote sensing technologies such as drones and satellites, remote sensing is now considered a fast, large-scale, and non-destructive method of data collection in the field of agriculture [1–3]. In the analysis of remote sensing images, vegetation indices (VI) are common features [4,5]. VI combines multiple bands from the sensor to estimate the vegetation vitality within a certain area, and is frequently used for plant classification, phenotyping, or monitoring plant-related parameters [4–6]. In the VI-based phenotyping protocols, the most common data processing involves the following steps: 1) obtaining images through remote sensing, 2) creating ortho-image of the studied field, 3) converting the images into VI, 4) cutting out each plot of interest from the ortho-images, and 5) averaging the VI values of all pixels in the plot to obtain a mean VI value at the plant level [4,7–11]. In parallel, the target phenotypic values (PV) of plots in the studied field are measured to correlate them with the average VI. Finally, this correlation is used to predict the trait values of all plots in the studied field [2,9,10]. However, VI-based phenotyping following this procedure frequently shows that the developmental stage of crops affects the performance of PV estimation [3,9,10]. For example, when predicting the yield of winter wheat via multispectral drone images, the testing R^2^ obtained from heading stage imagery was up to 0.56, which was 3.2 times higher than using images from the ripening stage [9]. Similarly, using images to predict rainfed corn yield around the silking stage (R1) is threefold better than the dent stage (R5) [10].

In the two examples presented above, researchers attributed these differences among growth stages to the differences in leaf area size and the differences in strength of plant physiological responses [9,10]. However, we hypothesize that another source of bias exists in the data processing. Frequently, there exists a non-linear relationship between VI and PV [6,8,12]. For example, up to 65% and 74% of the studies use non-linear functions, such as polynomial, logarithmic, and exponential curves, to predict the leaf area index (LAI) of maize and wheat [6]. When predicting the plant growth stages or growth-related phenotype by VI, the function between VI and phenotype would be an S-shaped curve [12]. Since the relationship between VI and PV, referred to as the link function, is frequently non-linear, averaging VI of all plants in a plot to obtain VI at the plot level will introduce bias. This is the so-called Jensen’s inequality [13], also referred to as ‘*the fallacy of the average*.*’* It states that for a nonlinear function *f(x)*, the average of it 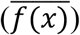 is not equal to the function of average *x*, which is 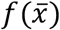 (Eq. 1) (14).

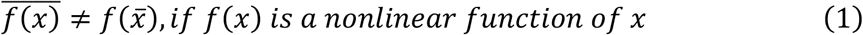

This fallacy of the averages only disappears if the link function *f(x)* is a linear function of VI. In that case, linearity would ensure that the order of taking the average or into the function does not matter [14].

Moreover, the degree of nonlinearity in a link function can vary across different ranges of VI values, and the VI distribution within a plot influences its range, mean, and variability. Consequently, the within-plot distribution of VI introduces an additional dimension to Jensen’s inequality. According to the results of previous studies, both the mean and shape of the VI distribution change with plant phenology [4,15,16]. For example, the normalized difference vegetation index (NDVI), one of the most common VI, would gradually increase with maize growth during the vegetative stage and decrease in the reproductive stage but the coefficient of variation of NDVI reaches the highest values in the middle of the vegetative stage, decrease to the lowest values at the beginning of the reproductive stage, and increase again at the harvest [16]. The distribution of VI changes during plant growth, and due to the non-linear property of the link function, averaging VI of each plot under different phenological stages introduces varying degrees of bias. This bias leads to the differences in performance observed under different phenological stages and finally constrains the performance of the PV prediction.

In this study, we first hypothesized that the strength of bias is affected by the type of link functions and distribution of VI. To demonstrate the extent to which the bias in using the average VI of each plot to predict the corresponding phenotypic values, statistical simulations with different link functions and VI distributions were used in combination with a real-world data dataset. To understand how to minimize this bias, we proposed a correction method that predicts the trait value before averaging to eliminate non-linearity and quantifies the potential improvement under different link functions, distributions, and measurement noise strengths. Based on our findings, we provide a guideline for researchers to enhance the image data processing procedure.

## METHODS

### Basic assumptions of the simulation studies

In all simulations, each experimental unit (in the context of a field experiment, an experimental unit equal to a plot) was assumed to contain 1,000 plants, with each plant corresponding to one pixel. For plant i in a plot, a vegetation index (*VI*_*i*_) and a phenotypic value (*PV*_*i*_) were assigned. The VI values for all plants within a plot were independent and identically distributed (i.i.d.), followed a beta distribution, a continuous distribution bounded between 0 and 1 (Eq. 2), chosen to mimic the typical range of NDVI.

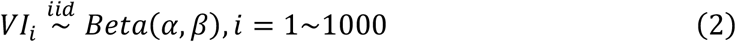

The probability density function (pdf) of the beta distribution can be adjusted by varying the parameters *α* and *β* (Eq. 3), enabling representation of a wide variety of shapes—including J-shaped, positively or negatively skewed, and approximately bell-shaped forms [17].

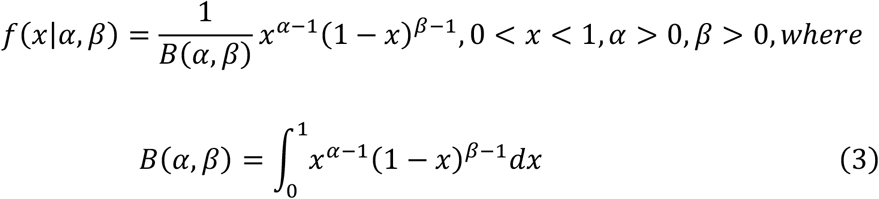

### Framework of simulations

We identified three factors that influence the effect of estimating PV at the plot level. The first factor is the effects of VI distribution on the bias of estimating PV. The second factor is the effects of the non-linear relationship (link function) between VI and PV on the bias of estimating PV. The first and second factors together are the two sources that determine the degree of fallacy in the average. The third factor is the effect of measurement noise, both from VI and PV in the training dataset, on predicting PV.

To test the effect of these three factors, we conceptualized a series of simulation studies. Each simulation involved: (1) defining the VI distribution, link function parameters, and noise level; (2) sampling VI values; (3) generating PV via the specified link function; (4) adding noise; (5) predicting PV; (6) averaging within plots; and (7-9) quantifying bias. The workflow was illustrated in Fig. 1. All simulations were conducted in the R environment [18].

**Figure 1.**
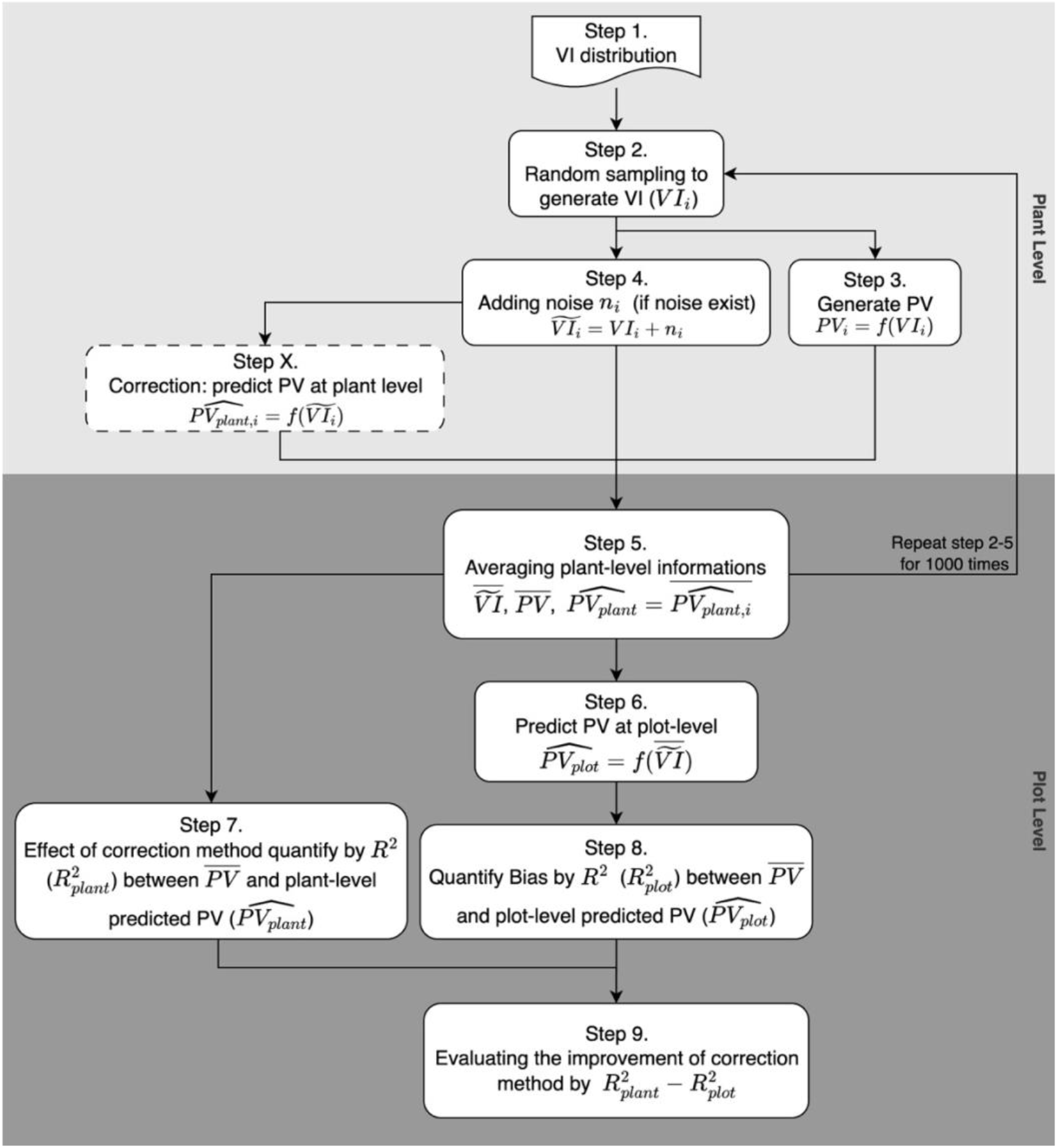
Framework of the simulation workflow. Vegetation index (VI) values were randomly sampled from the given VI distribution (Steps 1 and 2), where subscript *i* denotes the value from the i^th^ plant in a plot (*i* ranges from 1 to 1000). Phenotypic values (PV) were obtained by applying the link function to each VI (Step 3). When applicable, noise (*n*) was added to VI to produce noisy values, denoted as 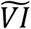, which represented the VI with noise added (Step 4). An additional step, predict PV at the plant level, was performed by obtaining the plant-level predicted VI 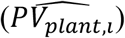 by applying the 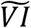 with the link function. All the values for each plant were averaged separately to obtain the mean VI 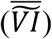, mean PV 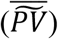 and mean predicted PV 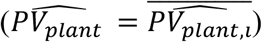 (Step 5). Steps 2–5 were repeated 1,000 times to generate results for 1,000 plots. Mean VI for each plot was used to predict PV at the plot level 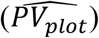 via the link functions (Step 6). The coefficient of determination (R^2^) between the mean PV and the two predicted PV from different levels were calculated to quantify the effect of the correction method and the bias (Steps 7 and 8), and the improvement of the correction method was evaluated by the R^2^ from the correction method minus the R^2^ from the conventional method (Step 9).

### Creating VI distribution

The simulation processes started with creating different distributions of VI (Step 1 in Fig. 1) through parameterization of beta distributions (Eq. 3), then conducting random sampling for 1,000 samples from the distribution. We used two different series of simulations to represent, and each of them contains a series of distribution parameter settings (Fig. 2).

**Figure 2.**
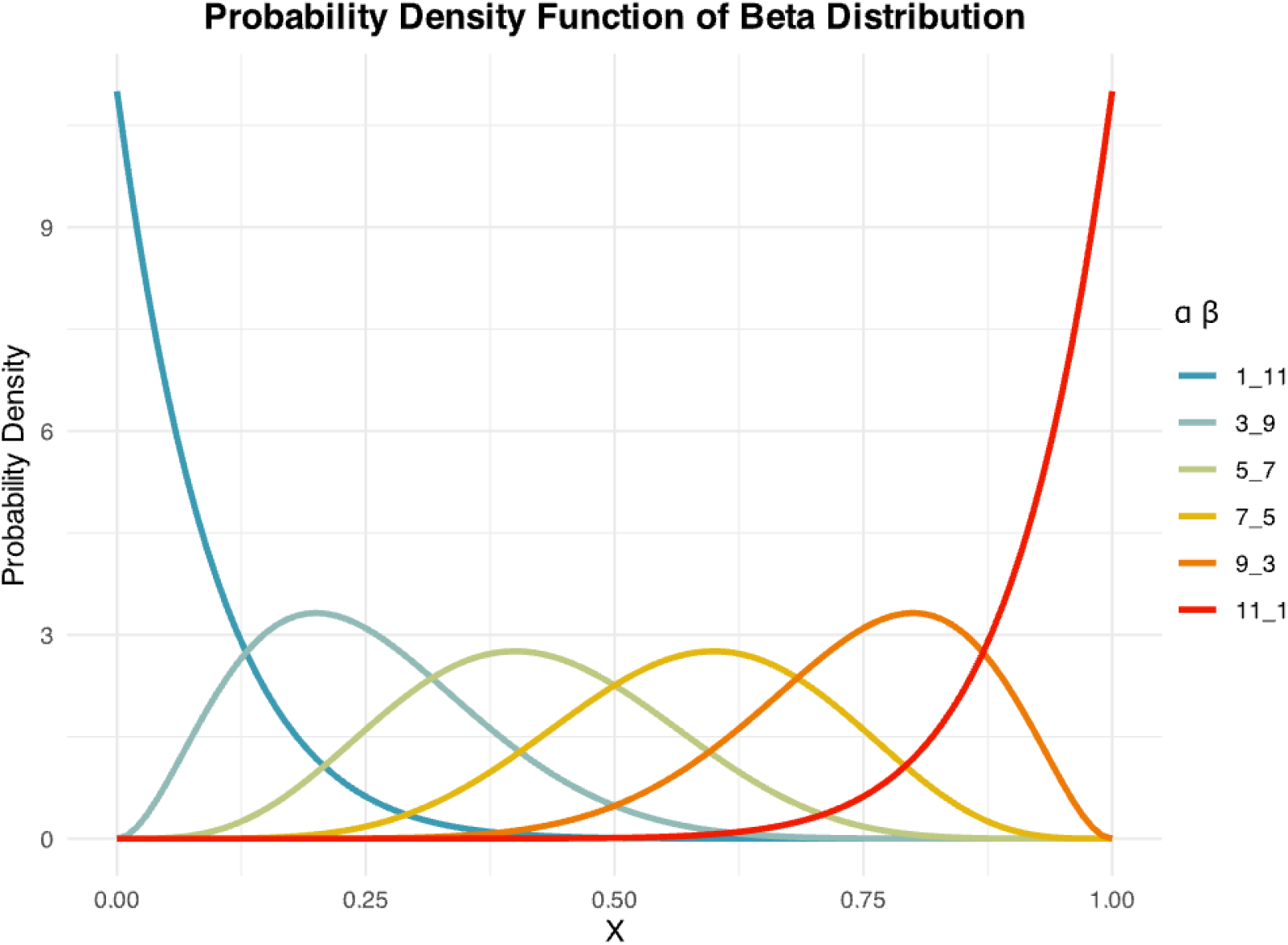
Probability density function of the six beta distributions used in the simulations. Each line color corresponds to a specific parameter pair (*α* and *β*) with values shown before and after the underscore, respectively.

The first series of simulations used the theoretical distributions based on the characteristics of the beta distribution. Based on Jensen’s inequality, the nonlinearity (also known as ‘curvature’) is the key factors that affect the strength of bias [14]. Since the nonlinearity of link functions is inconsistent in each local region of VI, the mean, skewness, and other statistical properties of the statistical distribution behind VI also impact the magnitude of bias. To capture these effects, we generated six beta distributions with parameters *α*= {1, 3, 5, 7, 9, 11}, and *β* = 12 – *α*. The sum of *α* and *β* was fixed to maintain a consistent central tendency while varying the mean and skewness. These distributions ranged from strongly right-skewed to strongly left-skewed, allowing assessment of bias under both realistic and extreme scenarios (Fig. 2).

The second series of simulations was designed to understand to which the fallacy of the average biases the estimation of phenotypic values in a real-world experiment. We used the NDVI images obtained from a paddy rice field experiment (unpublished) to create the distribution of VI. This experiment was conducted in New Taipei City, Taiwan, from March to July 2024. The multispectral images were collected at an altitude of 30 meters by a multirotor drone with a Micasense RedEdge MX multispectral camera on March 25 (transplanting), May 10 (tillering stage, 46 days after transplanting, DAT), June 7 (heading stage, 72 DAT), and July 11 (before harvesting, 105 DAT). These images were radiometric calibrated and used to create the orthomosaic of each band and converted into NDVI by Pix4D mapper [19]. Images of experimental fields were clipped, and soil pixels were removed before extracting NDVI values of rice plants. We estimated the parameters (*α* and *β*) of beta distributions at four imaging dates by maximum likelihood estimation by the R package *fitdistrplus* [20]. Parameters for intermediate dates without an image were obtained through interpolation.

### Link function transforming

After generating VI values, we converted them into PV by link functions (Step 3 in Fig. 1). Two types of link functions were used in this study (Eq. 4, 5).

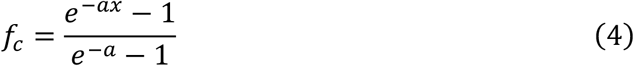

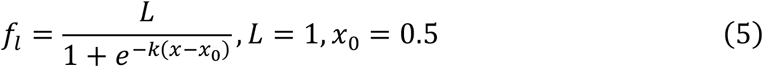

The first function, *f*_*c*_, is a monotonically convex/concave function (Fig. 3a). The shape of *f*_*c*_ is controlled by the parameter *a* to show the property from concave (which looks like a log function) to convex (which looks like an exponential function). Six levels of parameters, *a =* {-10, -5, -3, 3, 5, 10}, were used to represent different concave/convex link functions. Greater |*a*| values indicate stronger nonlinearity. The second function, *f*_*l*_, is a logistic function that is an S-shaped curve and was used in many growth-related models (Fig. 3b). Parameters *L* and *x*_*0*_ were fixed as 1 and 0.5 to center the inflection point. The parameter *k* was varied at six levels, *k =* {1, 3, 5, 10, 15, 20}, to demonstrate the effect of steepness. The nonlinearity of *f*_*l*_ increased when *k* increased and turned into a linear function when *k* = 1.

**Figure 3.**
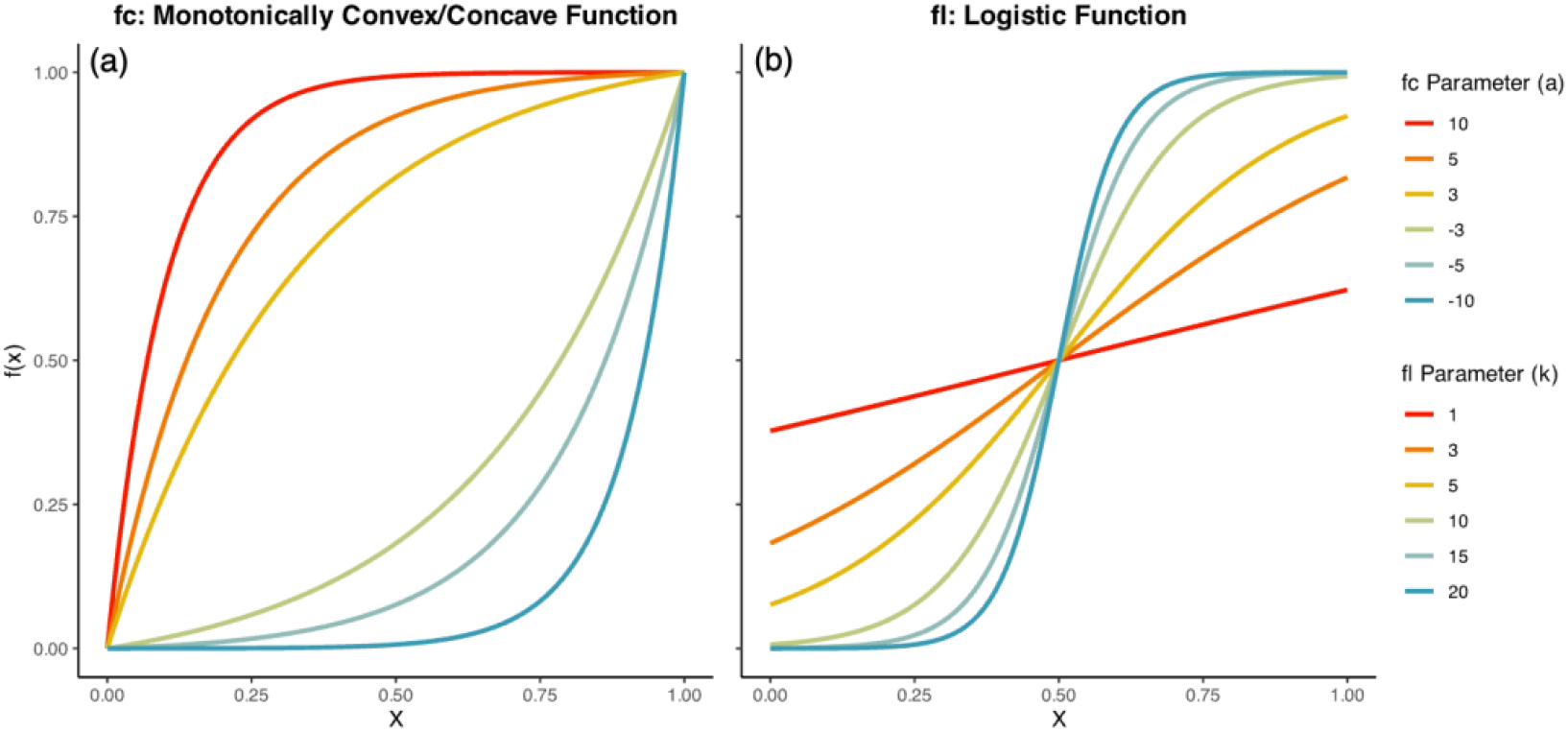
Link functions used in simulations. Two types of link functions, convex/concave function (*f*_*c*_, a) and logistic function (*f*_*l*_, b), were employed in this study to simulate different relationships between vegetation index and phenotypic value (Step 3 in Fig. 1). For each type of link function (parameter *a* for *f*_*c*_ and *k* for *f*_*l*_), six parameter levels were configured to evaluate responses across varying degrees of nonlinearity.

### Measurement noise setting

To examine how measurement noise affects PV estimation, random noise (*n*_*i*_) was added to the VI values (Step 4 in Fig. 1). This noise was sampled from a normal distribution, with its intensity controlled by a noise-level parameter *l*. The standard deviation of the noise distribution was defined as *l* multiplied by the standard deviation of the VI distribution. Increasing *l* increased the variability of the noise distribution, thereby strengthening the noise (Eq. 6). Simulations were conducted under four conditions: without noise adding, and noise levels *l* = {0.5, 1, 2} to assess the resulting performance changes.

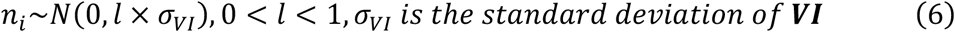

### Correct the fallacy of the average by predicting PV at the plot level

Since the non-linearity of link functions causes the fallacy of the averages, we proposed a correction method to eliminate the bias. Instead of the conventional approach—calculating the mean VI of each plot and then predicting PV at the plot level (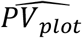, Step 6 in Fig. 1) —our proposed method reverses the order of operations. First, the VI for each plant was converted to a predicted PV 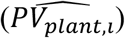 by the specified link function at the plant level (Step X in Fig. 1). This transformation linearizes the relationship between PV and 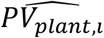, ensuring that subsequent averaging does not reintroduce bias. Since the noise was added before the plant-level prediction, (if noise was added), the effects of noise were also shaped by the nonlinear link function. The plant-level predicted PV values 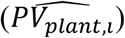 were then averaged to obtain the mean plant-level predicted PV for each plot 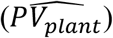.

We compared this corrected approach with the conventional method by calculating the coefficient of determination (R^2^) between the mean observed PV per plot and both 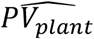 (corrected) and 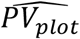 (conventional). This comparison was performed across all combinations of VI distributions, link functions, and noise levels to evaluate the effectiveness of the correction method.

### An interactive simulation application based on R Shiny

To facilitate the application of our simulation framework to diverse experimental conditions, we turned the whole simulation process into an interactive R Shiny web application. Users can upload their daily distribution parameters obtained from their experimental data (same as the second series of simulations), and set up other simulation settings (link function type, parameters, and noise levels), to verify the simulation results under the given conditions.

## RESULTS

### The fallacy of the average reduces the validity of using vegetative indices to predict phenotypic values by up to 82%

In our simulation study (Fig. 1), we used the coefficient of determination (R^2^) between mean PV and plot-level predicted PV for each plot to quantify the fallacy of the average (Step 8 in Fig. 1). Using a set of theoretical distributions without noise (Fig. 2), the R^2^ varies under different combinations of link function parameters and distribution parameters. When link functions were linear or nearly linear (*a* = 3 or -3, *k* = 3 or 5), R^2^ approached or equal to 1 (Fig. 4). Increasing nonlinearity in the link functions led to a reduction in R^2^, with the magnitude of this reduction depending on the distribution of VI. The most pronounced decline occurred with a logistic link function (*k* = 20) and distribution parameters *α*= 1 and *β* = 11, where R^2^ decreased to 0.18, 82% lower than the linear cases. Overall, we clearly demonstrated that greater linearity in the link function was associated with a smaller bias.

**Figure 4.**
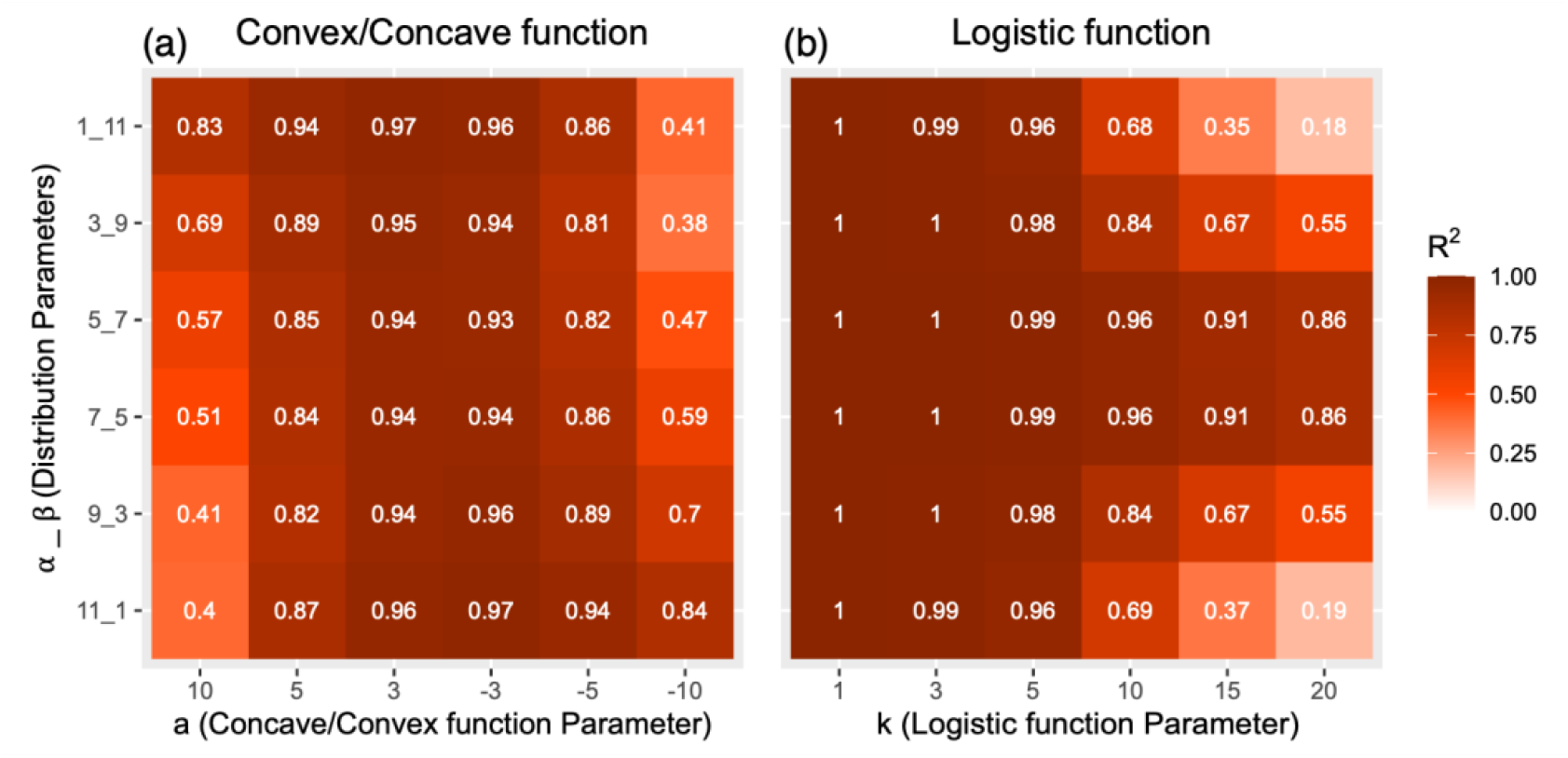
Correlation coefficient (R^2^) between simulated phenotypic values (PV) and the predicted PV at the plot level 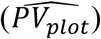. VI distributions within plots were simulated using beta distributions with varying shape parameters (*α* and *β*, Step 1 in Fig. 1). For each simulation, pixel-level VI values within a plot were transformed into PV using either (a) concave/convex functions or (b) logistic functions (step 3 in Fig. 1). No measurement noise was added in the simulation (*n*_*i*_ = 0, step 4 in Fig. 1). The VI for all plants in the plot were first averaged than used to predict 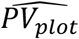 (Step 5 and 6 in Fig. 1), and the R^2^ between PV and 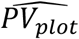 were calculated (Step 8 in Fig. 1). The degree of non-linearity was controlled by parameters *a* (for concave/convex functions) and *k* (for logistic functions).

To showcase the impact of phenology-dependent VI-distribution on the performance of estimating PV, we conduct simulations by using the distribution of NDVI images obtained from a paddy rice experiment as input (Fig. 5a), and the shape of VI distribution under each DAT shown in Fig. S1. The R^2^ between mean PV and predicted PV at the plot level (Fig. 5c and d) was low to 0.42 and 0.73 throughout the growth period under convex/concave and logistic link functions, respectively. Similar to the theoretical simulation (Fig. 4), increasing linearity improved R^2^ regardless of the VI distributions (e.g., *a* = 3 or -3 in Fig. 5c, and *k* = 1 in Fig. 5d). Also, R^2^ decreased the degree of nonlinearity of the link functions. The lowest R^2^ under the concave/convex and logistic link functions occurred at DAT 104 and DAT 51, which were around harvesting and tillering. These results highlight that phenology itself can influence phenotyping accuracy through its effect on the distribution of VI. Depending on link functions and phenological stage, such as heading or harvesting, phenotyping performance may be reduced by up to 58%.

**Figure 5.**
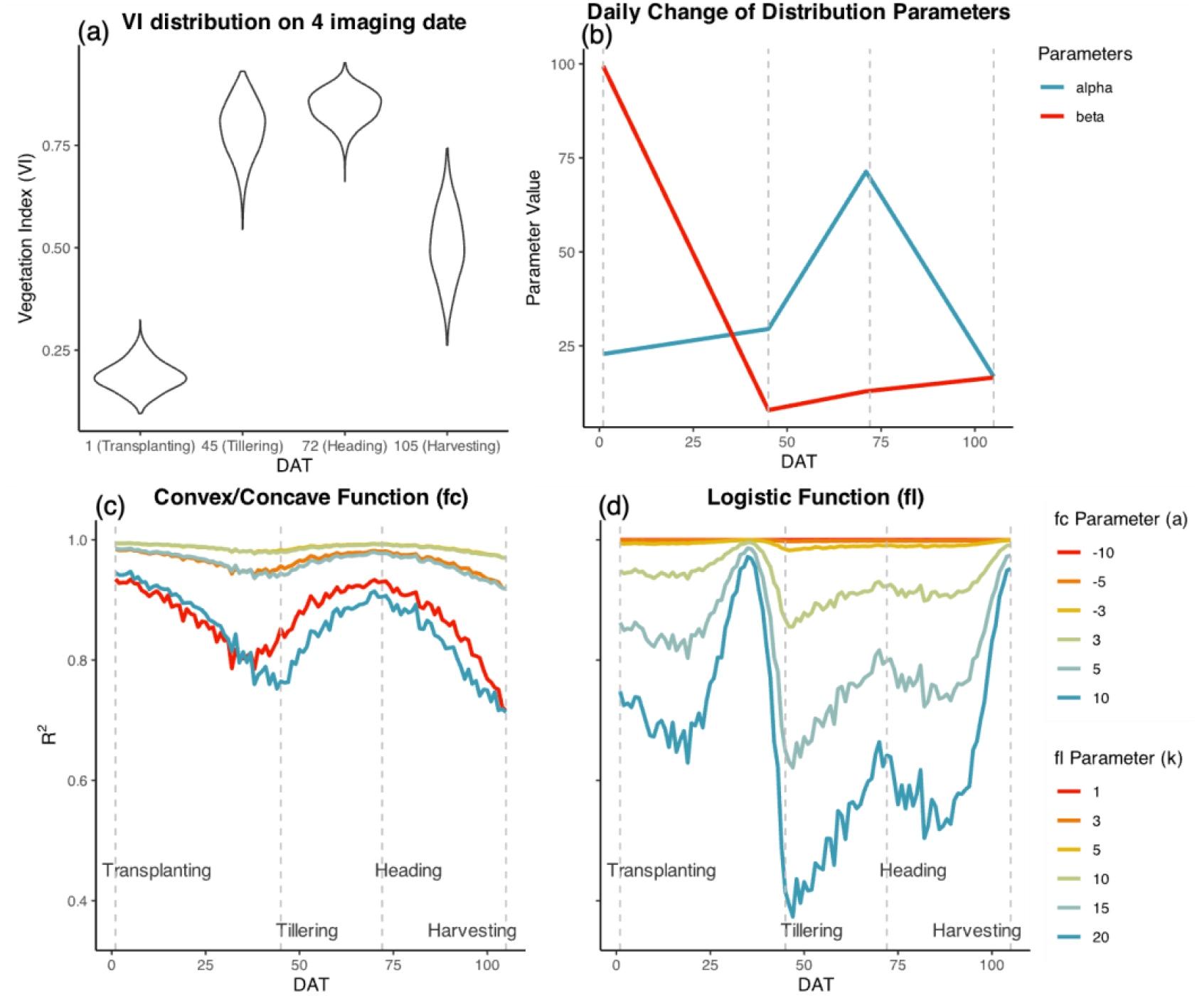
Impact of phenological stages on the correlation coefficient (R^2^) between simulated phenotypic values (PV) and the predicted PV at the plot level 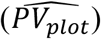. (a) The distribution of VI within plots was derived from the NDVI image of the paddy rice field experiment at the four imaging dates. The distribution parameters *α* and *β* were estimated from the distribution in (a) and interpolated between imaging dates (b). R^2^ between PV and 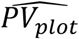 (c, d) of each day after transplanting (DAT) was simulated based on various parameter values for link functions and simulation setting shown in Fig. 3.

### Predicted trait before average can correct the bias arising from the fallacy of the averages

To assess the potential of the correction methods, predicted PV at the plant level before average, in mitigating the fallacy of the averages, we conducted a series of simulations under varying distribution types, link functions, and noise levels. Under the theoretical distributions in the absence of noise (Fig. 6), the correction method restored perfect linearity (R^2^ = 1.0, Fig. 6c and d, without noise), demonstrating that predicted PV before average can fully eliminate the biases introduced by averaging. However, once noise was introduced, the R^2^ values declined substantially—approaching 0.06 or 0.03 under the highest noise levels (Fig. 6a and b), indicating that measurement noise significantly impairs the prediction performance. With correction (Step X. Fig. 1), the R^2^ only reached a maximum of 0.61 under highest noise level (Fig. 6d, noise level = 2, *a* = 10, *α*= 1, *β* = 11), indicating that the performance losses caused by noise couldn’t be resolved by correcting the fallacy of average.

**Figure 6.**
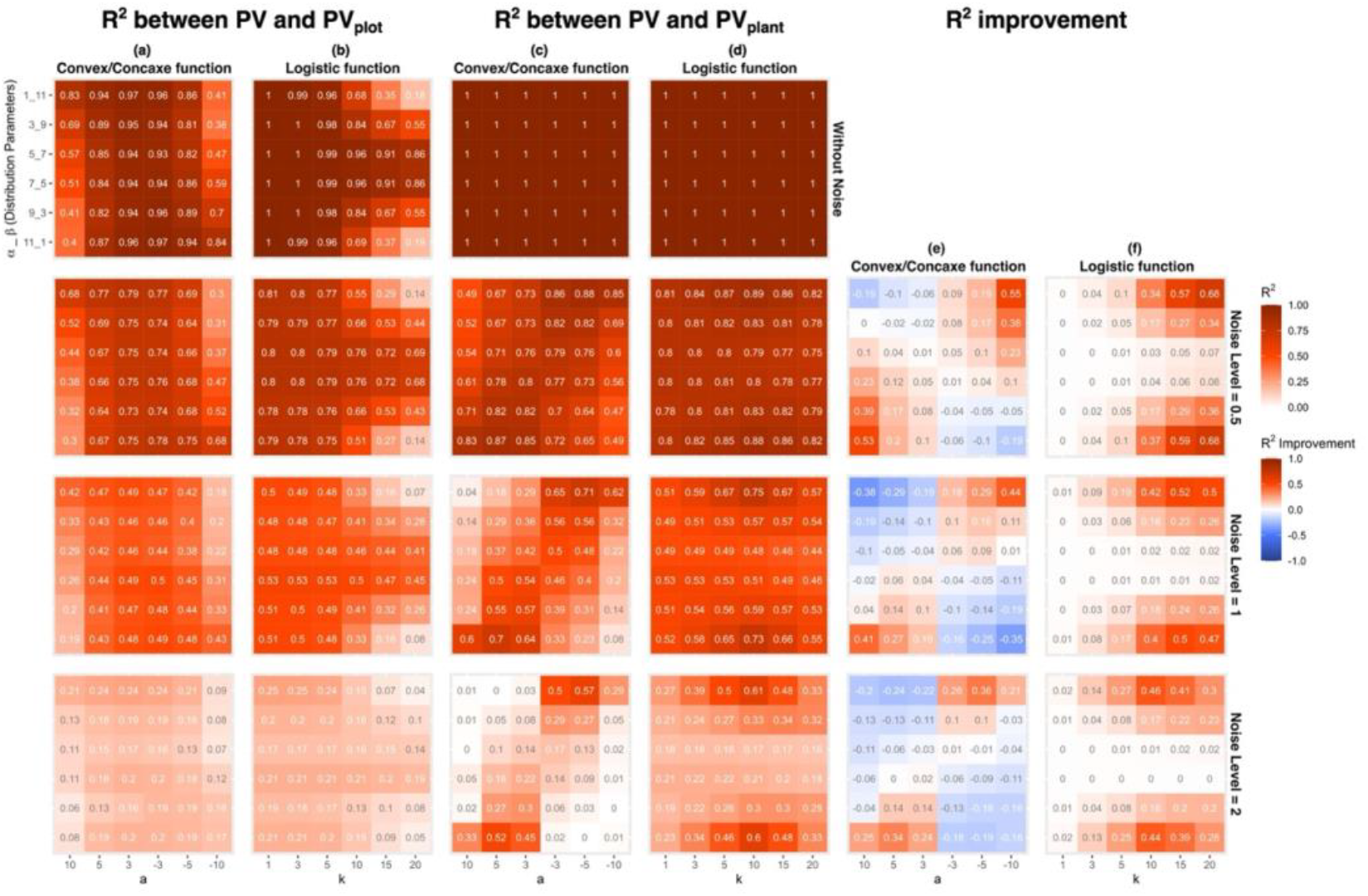
Correlation and potential improvement of bias between simulated phenotypic values (PV) and the predicted PV under different noise levels. The correlation coefficients were simulated based on different VI distributions, link function types, and parameter values of link functions. The simulation conditions used are shown in Figure 4. We conducted the PV prediction at both plot level (after average, step 6 in Fig. 1, a-b) and plant level (before average, step X in Fig. 1, c-d) to observe the improvement of the correction method (e-f) under different scenarios. Different levels of measurement noise were added in the simulation (noise level = 0.5, 1, and 2, step 4 in Fig. 1).

For convex and concave link functions, the benefit of the correction method was most evident along the diagonal axis of the heatmap (Fig. 6e). This pattern suggests that the correction method was most effective when the shape of the link function aligns with the skewness of the input distribution. For example, when concave functions were applied to left-skewed distributions (*α*= 11, *β* = 1, *a* = 10) or convex functions to right-skewed distributions (*α*= 1, *β* = 11, *a* = –10), the magnitude of improvement were 0.53 and 0.55 under the lowest noise level, respectively. Conversely, when the link function and the distribution skewness are misaligned, correction had a detrimental effect on model performance (e.g., *α*= 1, *β* = 11, *a* = 10; and *α*= 11, *β* = 1, *a* = –10). A consistent trend across simulations was the strong influence of noise on the effectiveness of the correction method. As noise levels increased, the benefits of correction diminished, and in some cases, became negative—especially in scenarios where the improvements at low noise levels were modest. Furthermore, the parameter combinations associated with the greatest improvements shifted with increasing noise: from those involving strong nonlinearity (|*a*| = 10) under low noise conditions (noise levels = 0.5 and 1), to moderate nonlinearity (|*a*| = 5) under higher noise (noise level = 2). These findings indicate that noise not only reduces the effectiveness but also alters the pattern of the outcomes of the correction method.

A similar trend was observed when applying the logistic link function (Fig. 6f). The effectiveness of the correction method varied with the interaction between nonlinearity and distribution skewness. The greatest improvement was up to 0.68, which occurred when strong nonlinearity coincided with highly skewed distributions, whether left- or right-skewed (e.g., noise level = 0.5, *α*= 1, *β* = 11, *k* = 20). Although noise reduced the maximum achievable improvement, all scenarios under the logistic link function showed positive improvements—there were no cases in which correction led to a reduction in R^2^.

Collectively, these simulation results emphasize three key insights. First, measurement noise is a critical factor influencing the predictive accuracy of VIs when used to predict phenotypic traits. Second, link function correction improves prediction performance across a wide range of conditions. Third, the magnitude of improvement by the correction method is contingent upon the interaction between the noise level, the VI distribution, and the link function.

Simulations based on paddy rice NDVI imagery (Fig. 7) revealed trends consistent with those observed in the simulations using theoretical distributions. Under noise-free conditions, the correction method fully restored perfect linearity (R^2^ = 1.0), reaffirming its capacity to eliminate the fallacy of the averages (Fig. 7c and d, without noise). Once noise was introduced, predictive performance declined before correction across all link function types, highlighting the disruptive effect of noise on the correlation. Even when using the correction method, the highest R^2^ achieved was only 0.35 under the highest noise level (Fig. 7d, *k* = 15, DAT = 45), once again demonstrating the severe damage caused by noise.

**Figure 7.**
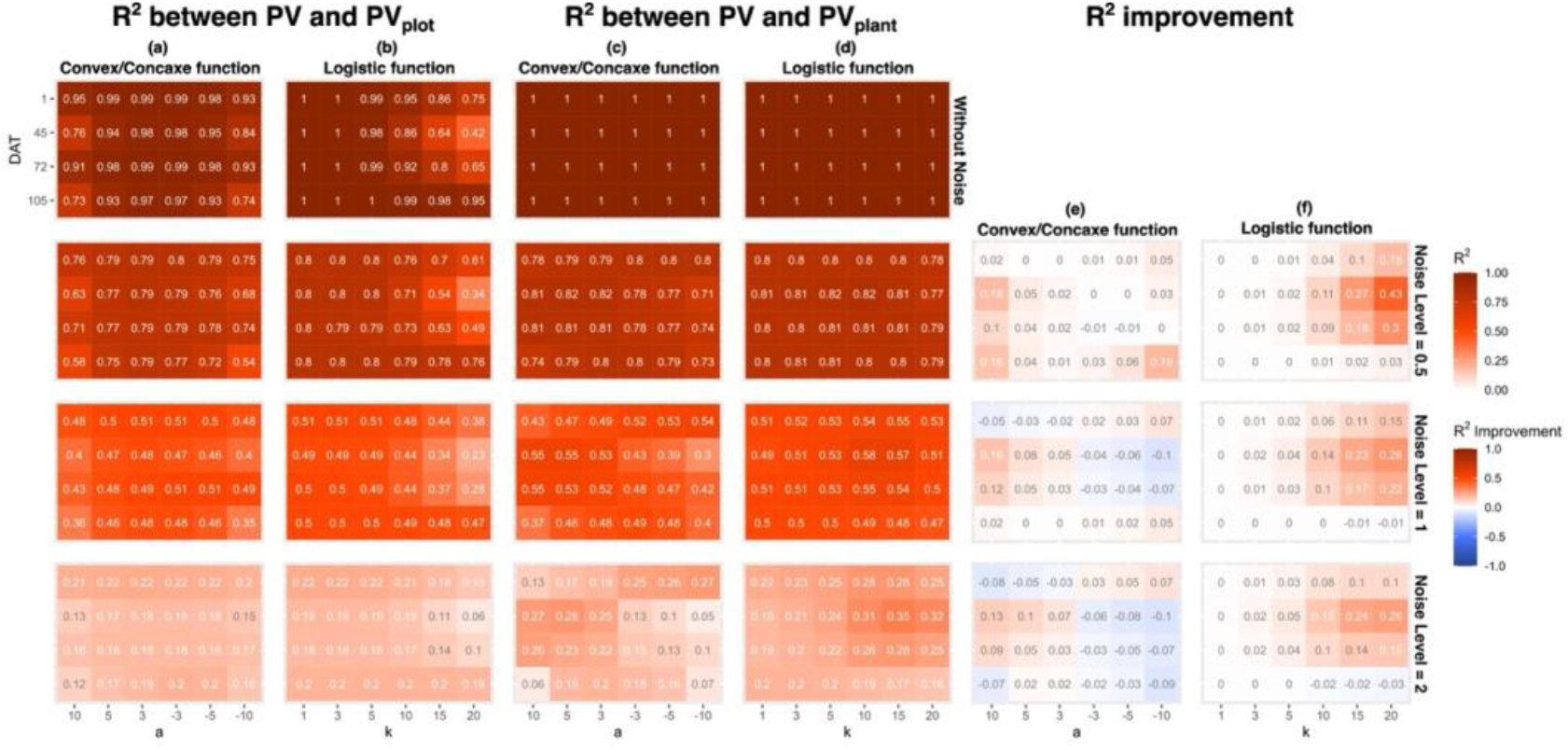
Correlation and potential improvement of bias between simulated phenotypic values (PV) and the predicted PV in real experimental data with different levels of noise added. The correlation coefficients were simulated based on vegetation index (NDVI) distributions derived from the NDVI images of a paddy rice experiment (Fig. 5a). The simulation condition setup was shown in Fig. 6.

As observed with the theoretical distributions, the magnitude of improvement after correction was influenced by the noise level. When using convex/concave link functions, the maximum improvement reaches 0.19, occurring when the lowest noise was added, around the harvesting stage, and under the large nonlinearity link function (Fig. 7e, noise level = 0.5, DAT = 105, *a* = -10). The correction improvement was also high (0.18) when around the tillering stage, lowest noise level, and large nonlinearity link function (Fig. 7e, noise level = 0.5, DAT = 45, *a* = 10). Conversely, in scenarios where correction provided only a small improvement under low noise, those improvements diminished or turned into a negative effect as noise levels increased.

For simulations using the logistic link function, the noise level also shows the impact on the improvement. Most of the scenarios exhibited the positive improvement after correction, with a maximum R^2^ increase of 0.43 at the tillering stage (Fig. 7f, noise level = 0.5, DAT = 45, *k* = 20). Notably, the largest improvements were consistently observed during the tillering stages, particularly when combined with link functions exhibiting higher nonlinearity (e.g., Fig. 7e, *a* = 10; Fig. 7f, *k* = 20).

Overall, these simulations—whether based on theoretical constructs or empirical NDVI data—underscore two critical point. 1) The measurement noise plays a pivotal role in determining predictive performance. 2) The correction method (predict PV before average) is effective in mitigating the fallacy of the averages, especially under conditions of low noise and strong nonlinearity.

## DISCUSSION

### High-throughput phenotyping should consider the fallacy of the average

According to Jensen’s inequality, the fallacy of the average occurs if 1) the relationship between image indices and trait values is nonlinear and 2) simply averaging all VI values in a plot without considering the distribution of VI within a plot [13,14,21]. Depending on the degree of nonlinearity in the link function and the shape of the VI distribution, the bias may range from zero to as high as 82% (Fig. 4), or even much higher if the nonlinearity keeps increasing. Since the link functions are nonlinear and the degree of nonlinearity varies across different VI ranges, the location of VI values concentration strongly influences the magnitude of the fallacy of the averages (Fig. 8). When VI values are concentrated in the more linear portion of the link function, the fallacy of the averages was lower (Fig. 8b). In contrast, when VI values are concentrated in the curved or nearly horizontal sections, the fallacy of the averages has a much stronger effect (Fig. 8c and d). In addition to the correction method, research in other fields has pointed out that using the statistics of the input variable’s distribution and other parameters that may affect the link function as input values for machine learning models may reduce bias [21]. However, the information of the VI (or the input variable) distribution and the link function is still necessary to minimize the fallacy of the averages during analysis.

**Figure 8.**
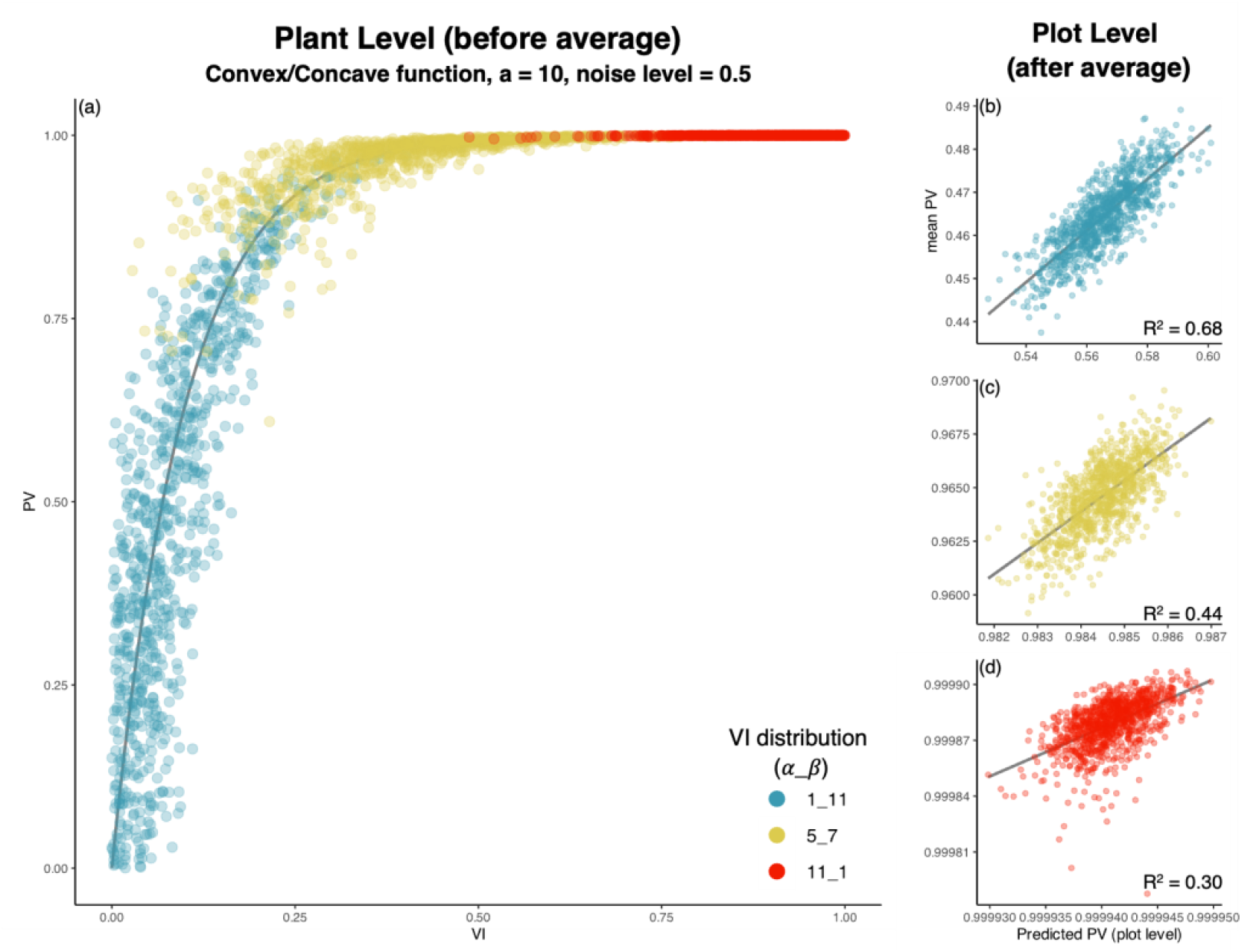
Examples showing how the interplay between distribution and concentration of the vegetation index (VI) values and the nonlinearity of link functions influences the performance of phenotyping (simulated R^2^). VI for each plant were sampling from three different VI distributions (*α* and *β* = 1&11, 5&7, 11&1, Step 1 in Fig. 1), and transformed to the phenotypic value (PV) by a convex/concave function with *a* = 10 (Step 3 in Fig. 1) under a noise level of 0.5 (Step 4 in Fig. 1) (a). For all three VI distributions (b-d), the step from sampling to averaging was repeated 1,000 times to obtain the value (Fig. 1, steps 2-5), and the mean VI for each plot was used to obtain the predicted PV at the plot level (Fig. 1, step 6). The value of the simulation R^2^ came from the simulation results in Fig. 6.

The shape of the VI distribution depends on the VI used, the crop type, and the phenology. Most VI follow a unimodal distribution [22], and the center of the distribution is influenced by phenology [16,23]. VIs established in the form of normalized differences (e.g., NDVI) are asymmetric, and their skewness depends on the correlation between the two bands in the combination [22]. Therefore, different VIs, or the same VI obtained from various crops, will affect the shape of VI distributions. In an experiment to calculate the RGB image-based VI for forage grasses, the results also showed that the trends in different indices over time varied [4]. However, the distributions of VI were not reported in most of the studies, and therefore, it became difficult to estimate the magnitude of the bias in their cases.

The type and nonlinearity of the link function depend on the target trait, the VI used, and crop species. For example, when using NDVI to predict wheat LAI, different studies propose different types of link functions, such as linear and stepwise linear [24,25], logarithmic curves [26,27], exponential curves [28,29], and power curves [30,31] under different vehicles, spatial and temporal scales. In studies that use VI to estimate or predict phenology, S-shaped curves similar to or the same as *f*_*l*_ were commonly used to fit the relationship between VI and phenology [12,32,33]. However, none of these studies have addressed the potential impact of the fallacy of the averages on the performance, indicating that systematic bias can substantially limit the power of high-throughput phenotyping.

Based on the two aspects mentioned above and our results, it can be concluded that in high-throughput phenotyping, the strength of the fallacy of the average’s impact on estimated results depends on crop type, target phenotype, the VI used, and phenology. This could be one of the reasons for the phenotyping performance differently across different phenological stages, which has mainly been interpreted as differences in the plant growth and the intensity of plant physiological responses across time [4,9,10]. Due to different combinations of traits and VIs, the trends in predictive performance may differ from those observed in our study.

It is worth noting that the fallacy of the average caused by averaging an entire area of an image not only occurs in VI-based phenotyping but is also valid in all image-based prediction, from cellular to canopy levels. For example, when using hyperspectral imagery to predict the severity of plant disease, a common approach is to capture an image of the entire leaf and average the pixel values across all regions [34,35]. However, the severity of disease can vary from one part of the leaf to another, and the relationship between disease severity and spectral reflectance is often nonlinear. These together satisfy the criteria for the fallacy of the averages. A similar issue arises in using chlorophyll fluorescence imaging to study plant physiological responses, where the mean value of the entire leaf is often used [36,37]. Here, too, the physiological response can differ across regions of the leaf, and its relationship with the measured chlorophyll fluorescence signal may not be linear, which once again fulfills the conditions creating the fallacy of the averages. Overall, we recommend that researchers should carefully consider whether averaging two variables with a nonlinear relationship could introduce bias and potentially lead to inaccurate conclusions of further data analyses (e.g., correlation of traits and genome-wide association studies).

### Image-based phenotyping should include noise estimation to ensure data quality

Our results indicate that noise is a major factor influencing the performance of VI-based phenotyping and also affects the efficacy of correcting the fallacy of the average by the correction method. It is essential to quantify the strength of noise during the analysis process. Noise in VI-based phenotyping originates from three main sources: 1) sensor-related noise from imaging devices, 2) random environmental noises (e.g., atmospheric effects, weather and temperature, soil reflectance, etc.), and 3) noise introduced during the phenotyping processes [38– 40].

The noise from sensors can be measured in advance by measuring their signal-to-noise ratio (SNR) [38,39]. SNR can vary significantly depending on the sensor type and exposure settings. Under well-setting exposure conditions, the SNR achieved with hyperspectral imaging ranges may be lower than the lowest noise level, or between the lowest to medium noise levels, depending on different wavelengths [41,42]. When using a high-precision multispectral camera for measurement, the noise level was even lower [43]. For thermal imaging, the error range of consumer-grade thermal cameras can reach as high as ±5%, compared with professional-grade cameras for about ±1% [44]. Although these values cannot be directly compared with the noise level in our simulations, they can still be used as a reference to understand the noise level from the camera when information about the object being measured and the measurement environment is available. At the same time, less noise can also be achieved by using different VI combinations [40]. Random noises can be reduced or eliminated through an appropriate experimental design. For example, in drone image collection, noise in image quality can be minimized through field layout and flight planning [45]. Appropriate correction procedures can also effectively reduce noise. In measurements taken with multiple sensors and over time, failure to perform corrections can introduce up to 40% noise into NDVI [46]. The correction of atmospheric errors can reduce the error of NDVI up to 11.94% [47]. Noise from phenotyping can be measured and limited by performing technical replicates [48]. Even though these noise measurements are not yet widely regarded as part of image-based phenotyping [40], we believe that future studies should more actively measure noise to understand the extent to which noise contributes to error in the final research results and how it can be considered in the context of correction for the fallacy of the average.

## CONCLUSIONS

This study aims to systematically investigate the fallacy and bias introduced by averaging VIs in plant phenotyping when a nonlinear relationship exists between VI and phenotypic traits. Using mean VIs for trait prediction can lead to substantial, non-trivial estimation bias, which reduces predictive performance. The magnitude of this bias depends on both the nonlinearity of the link function and the distribution of VI values within experimental plots. When VI values are concentrated in the more nonlinearity sections, the fallacy of the averages becomes stronger. Since the VI distribution changes during plant growth, the magnitude of bias is also strongly influenced by plant phenology.

A possible method to correct the bias is to predict trait values at the pixel level before averaging. The effect of correction depends on the noise between the relationship of VI and the trait. The correction can effectively eliminate bias in noise-free situations and significantly improve predictive accuracy under low to moderate noise levels. Nevertheless, when the noise increases, the effect of the correction reduces or even turns into a negative effect in some cases.

In most of the previous studies, the distribution of VI and the strength of noise were not reported. We contend that those crucial information related to bias strength should be reported in future research. Furthermore, where appropriate, bias correction should be regarded as an essential procedure to ensure the validity of the findings. Moreover, the findings suggest that the similar bias also affects other image-based metrics—such as hyperspectral or chlorophyll fluorescence imaging—if mean values are used as predictors for nonlinear traits when the nonlinear relationship between index and trait, underscoring the need for careful bias evaluation and correction in plant phenotyping and related fields.

## Supporting information

Figures S1. The shapes of vegetation index (VI) distributions (NDVI) derived from the paddy rice experiment on each day after transplanting (DAT).

## Competing interests

The authors declare that there is no conflict of interest regarding the publication of this article.

## Funding

Chun-Han Lee was supported by the DAAD/NSTC Sandwich Scholarship Programme for providing the opportunity and funding to visit Germany and conduct this research. Li-yu Daisy Liu was funded by the Ministry of Agriculture, project number 114AS-1.2.3-AS-05. Tsu-Wei Chen was funded by Deutsche Forschungsgemeinschaft (German Research Foundation, DFG) under “Emmy Noether Programm”, project number 442020478.

## Author contributions

Chun-Han Lee: Methodology, Software, Formal analysis, Visualization, and Writing (Original Draft). Chih-Chen Tseng: Data Curation. Li-yu Daisy Liu: Data Curation and Funding acquisition. Tsu-Wei Chen: Conceptualization, Methodology, Resources, and Writing (Review & Editing)

## Data availability

The interactive R Shiny website is publicly available at https://71vuys-chun0han-lee.shinyapps.io/fallacy_of_average_shiny/. The paddy rice experiment images and simulation scripts will also be available on the website when the article is accepted.

## Notes

### Competing Interest Statement

The authors have declared no competing interest.

