## Supplementary figures and images for "The Fallacy and Bias of Averages on Vegetation Indices based Plant Phenotyping"

### Figures S1. The shapes of vegetation index (VI) distributions (NDVI) derived from the paddy rice experiment on each day after transplanting (DAT).

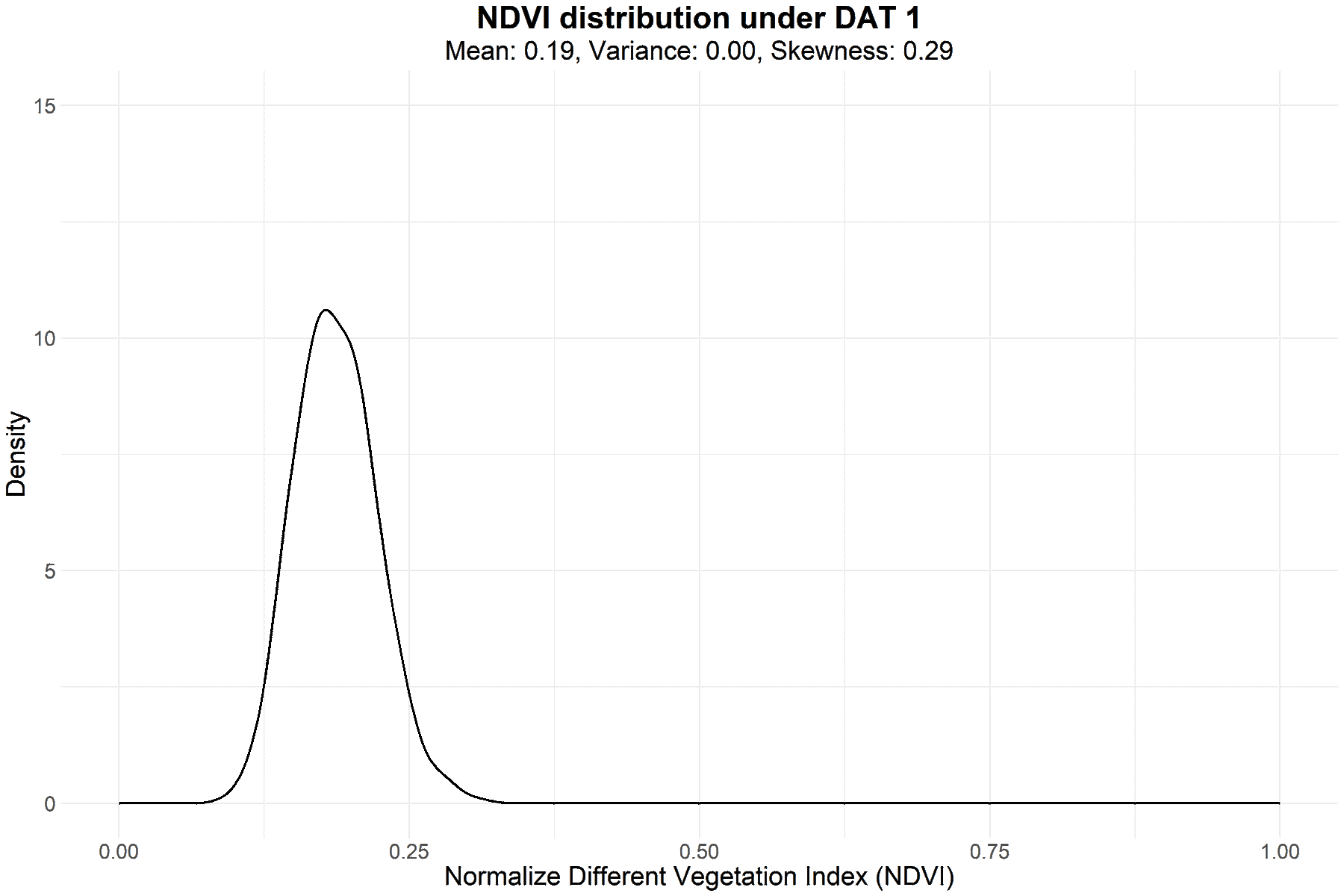
